# Directionally tuned signals in mouse visual and subicular cortex during passive rotation

**DOI:** 10.64898/2026.09.29.755383

**Authors:** Marin Dauguet, Flavia Aluisi, Mark Wexler, Ivan Cohen, Jean Laurens, Michael Graupner, Desdemona Fricker

## Abstract

Head-direction (HD) cells integrate visual and vestibular inputs and provide an internal compass which is essential for navigation, but how the directional activity in the subicular complex (SUB) and in the visual cortex (VIS) depends on the visual scene has not been compared directly. Here, we recorded simultaneously from the two regions with Neuropixels probes, while rotating head-fixed mice across all azimuthal directions, in controlled visual conditions, and in darkness. We found many directionally tuned units in both SUB and VIS. The preferred directions of SUB units were distributed uniformly and followed the rotation of a single prominent visual cue. The preferred directions of VIS units on the other hand were concentrated towards the visual cue. Heading was decoded with good accuracy from population activity from either region in cue rich visual conditions. In the presence of a single cue, decoding performance was higher in VIS. In darkness, the directional tuning was apparently lost at the single cell level, in both VIS and SUB. Importantly, even in darkness, the tuning-ordered structure of pairwise correlations was largely maintained in SUB, but not in VIS. This dissociation suggests that the SUB coherence reflects an internally organized attractor that can maintain its structure without vision or active movement, whereas VIS coherence largely reflects the external sensory structure. A wave-like distortion in decoded heading with respect to a visual cue suggests that a single salient landmark can shape the HD signal. Taken together, visual and head direction signals allow for an internal compass with continuity in the absence of landmarks, and recalibration as landmarks are reinstated.

## Introduction

The HD system integrates multimodal sensory inputs, including self-motion and landmark cues, to maintain and update an internal sense of direction (J. S. Taube 2007; Peyrache et al. 2015). However, how the directional activity depends on the visual scene when passively rotated remains poorly understood. Each HD cell fires selectively when the animal faces a particular direction, its preferred firing direction (PFD), but is independent of spatial position (J. S. Taube et al. 1990a, 1990b; J. S. Taube and Muller 1998). The population signal yields a consistent and stable HD signal that orients behaviour (Butler et al 2017). HD cells were first reported in the rat dorsal presubiculum, also termed postsubiculum (Ranck 1984), and quantitatively characterized there (J. S. Taube et al. 1990a, 1990b; J. S. Taube and Muller 1998). The presubiculum (PreS) is part of the subicular complex, located in the parahippocampal formation, a key area for the processing and integration of spatial and visual information. It receives directional inputs from the anterior dorsal thalamus (ADn; Yoder et al. 2011) and projects to the medial entorhinal cortex (MEC; Preston-Ferrer et al. 2016); a subpopulation of PreS neurons also projects back to the lateral mammillary nucleus (LMN; Huang et al. 2017). The PreS is thus well positioned to update the subcortical internal compass with external landmarks information (J. S. Taube and Bassett 2003).

Head direction signals are distributed along a series of pathways that ultimately produce a subject’s sense of orientation. LMN encodes rotational velocity by representing the angular velocity that accompanies the subject’s movements related to their head direction (R. W. Stackman and Taube 1998). ADn serves as a primary relay for the HD signals generated in these subcortical circuits to the cortex (J. S. Taube 1995; J. S. Taube 2007). The retrosplenial cortex (RSC) provides HD signals with allocentric reference frame navigational signals, reflecting stable visual landmarks (Wiener and Taube 2005; Vann et al. 2009). Cortical areas, including the primary visual cortex itself, carry head-motion signals of vestibular origin (Vélez-Fort et al. 2018), and angular head velocity is encoded multisensorially in retrosplenial circuits (Hennestad et al. 2021; Keshavarzi et al. 2022). HD signals in the MEC and parasubiculum (ParaS, also part of the subicular complex) are associated with grid cell and velocity signals, placing head direction within a navigational metric that organizes spatial cues into a unified internal map (Boccara et al, 2010; Spalla et al 2022; Sargolini et al. 2006; Moser et al. 2008). Here we focused on the subicular complex, and in particular on PreS, as a likely candidate for linking the internally generated direction estimates from ADn to visual landmark signals from RSC (Richevaux et al. 2025).

As the primary visual cortex carries head motion signals and also responds to visual landmarks, a simultaneous comparison of subicular complex and visual cortical populations can separate what each region contributes. The visual system provides critical information to anchor or reset the body’s internal compass (J. P. Goodridge and Touretzky 2000; Zugaro et al. 2003; Knight and Hayman 2014). Visual anchoring ensures that the ring attractor dynamics of the HD system remain correctly locked to the salient objects in the world (Skaggs et al. 1995; Redish et al. 1996) as has been demonstrated in classical cue shift experiments (J. Goodridge and Taube 1997; Zugaro et al. 2003). Without reliable visual input the internal compass will drift, leading to cumulative error (J. S. Taube et al. 1990a, 1990b; J. P. Goodridge et al. 1998; Knierim et al. 1998). Previous head-fixed studies have demonstrated presubicular HD tuning during passive rotation in the rat (Preston-Ferrer et al. 2016) and visual anchoring of heading signals in RSC in the mouse (Sit and Goard 2023). Building on these, we compared subicular and visual cortical responses to landmark manipulations while replaying the same rotation sequence across different visual conditions.

Computational modeling posits that coherence in the HD network arises through attractor dynamics, where neurons tuned to neighbouring head directions receive mutual excitation, while the inhibitory connections confine the spatial breadth of the firing to a localised region or “bump”. This bump of activity follows angular head velocity signals so that it slides smoothly as the animal turns its head (Sharp et al. 2001; Bassett and Taube 2001). Most of our knowledge about HD coding has come from freely moving animals. Those conditions are sensorily crowded because vestibular, proprioceptive, and motor efference signals vary at the same time. Landmark rotation and cue-conflict experiments have revealed how the system arbitrates among competing inputs (J. Goodridge and Taube 1997; Zugaro et al. 2003), but the specific computations of how visual, vestibular, and motor-related signals are combined remain unknown. A key prediction of attractor models is that the relationships of neurons active together should persist even when the external reference frame is lost, as observed across sleep and wake in ADn (Peyrache et al. 2015).

Although much progress has been made in understanding the dynamics of the HD system, major questions remain regarding how directional signals are generated and maintained when self-movement is absent. Passive rotation itself does not abolish HD firing in the ADn (Shinder and Taube 2011), but passive transport degrades the updating of the directional estimate by path integration (Stackman et al. 2003). The passive rotation of a head-fixed animal preserves vestibular signals, while minimizing motor, efference copy and proprioceptive signals. In such a paradigm, the stability of directional coding may depend on attractor dynamics that preserve internal coherence, on the visual signals that provide external alignment, or on their interaction. How simultaneously recorded subicular and visual populations differ in this respect across landmark manipulations and darkness has not been examined.

Here, we sought to determine how the visual scene shapes the directional activity in the subicular complex and in the primary visual cortex of head-fixed mice undergoing passive rotation. To this end, mice were secured on a rotating platform and exposed to the experimental room or a controlled visual surround with a single visual landmark. Neural activity was recorded with high-density Neuropixels probes across the pre-, post-, para- and subiculum (SUB) and also the visual cortex (VIS). We asked whether single unit directional tuning in each region follows the landmark, whether the population readouts of the two regions are anchored to it in the same way, and which properties of the directional code persist in the absence of landmarks.

## Methods

### Animals

Experiments were conducted on six mice (Mus musculus, four male, two female), aged between 4 and 5 weeks. The mice were sourced from locally sustained lines on C57BL/6 J background (1 VIP::Tom, 1 B6_PVcre; 4 wild-type) and housed in groups of 2-5 animals in ventilated cages under a controlled environment with a 12 hours light/dark cycle (lights on from 7:00 AM to 7:00 PM). Food and water were provided ad libitum. Animal care and handling adhered to the guidelines set forth by the European Community Council Directive 2010/63/EU on the protection of animals used for scientific purposes and complied with French legislation, related to the protection of animals used for experimentation (Ministère de l’Enseignement supérieur et de la Recherche). All procedures were reviewed and approved by the local Institutional Animal Care and Use Committee (C2EA-34) and by the French Ministry of Research 01025.02 (APAFIS #49200-2024021823341167 v7).

### Stereotaxic surgery

Mice underwent a preparatory surgery under general anesthesia, in which a head implant was fixed to the skull for head-fixation on the rotating stage and a craniotomy was opened at a site from which one probe penetration would cross the primary visual cortex and the underlying subicular complex. Anesthesia was induced with a 3% concentration of isoflurane, then reduced to 1% for maintenance. The depth of anesthesia was confirmed to be adequate based on the absence of the paw withdrawal reflex and the stabilization of breathing. The mice’s body temperature was maintained at 37°C using a thermostatically controlled heating pad to prevent hypothermia during surgery.

After each mouse was placed in a stereotaxic frame, the scalp was cleaned of hair and disinfected with 10% povidone iodine solution (Betadine), and lidocaine (4 mg/kg, subcutaneously) was injected into the scalp to block sensation of pain in the underlying tissue. A small unilateral craniotomy was placed with a 0.5 mm burr above the right visual cortex and subicular complex (AP −4.0 mm, ML +2.3 mm, relative to bregma; Paxinos and Franklin 2013). The craniotomy was then sealed using Kwik-Cast silicone sealant until the day of the recording. A customized titanium head implant (Titanium Grade 5, Ymetry, France) matching the curvature of the skull was then positioned over the exposed skull and bonded with dental cement (C&B Super-bond). The cement was applied carefully to ensure that it did not drip onto surrounding tissue and that a strong bond was formed between the cement and the cranial surface. Finally, the surgical site was flushed with sterile saline and cement was smoothed to minimize irritation to the skin. The opened skin surrounding the head implant was attached to the dental cement pedestal using veterinary tissue adhesive (VetBond).

Buprenorphine (0.05 mg/kg, subcutaneously) was administered as preoperative analgesia to alleviate post-surgical pain and to reduce the amount of stress experienced by the mice during the surgical procedure. Carprofen (20 mg/kg, subcutaneously) was administered as postoperative analgesia as a nonsteroidal anti-inflammatory drug. The mice were then placed in a warm recovery chamber until fully conscious, and were allowed a recovery period of at least 24 hours before subsequent manipulations. The recordings were performed between 1 and 2 days after the surgery.

### Extracellular recordings using Neuropixels

On the day of recording, the mouse was head-fixed at the centre of a motorized rotating stage (RGV160BL, Newport). The silicone seal was removed from the craniotomy, the dura was resected over the exposure, and the surface was kept moist with sterile saline. Extracellular spiking activity was recorded using Neuropixels probes 1.0 (Jun et al. 2017). Prior to insertion, each probe was coated with the fluorescent lipophilic dye DiI (Thermo Fisher Scientific) to facilitate post hoc histological verification of the probe track. The probes were inserted using a high precision four axis micromanipulator (Luigs & Neumann).

Target regions were the primary visual cortex, the postsubiculum and the presubiculum. The probe was advanced vertically, with no tilt in either the mediolateral or the anteroposterior axis, at 10 µm/s, to a final depth of 3 mm below the brain surface. The craniotomy was covered with sterile saline throughout insertion and recording. Neural activity was monitored in real time during insertion. Recordings began at least 15 minutes after the probe reached its final depth, to allow the tissue to settle.

Electrophysiological data were acquired using the SpikeGLX software package (Janelia Research Campus, Bill Karsh, 2023). Recordings used the 384 channels of the first bank, at the probe tip. Signals were split into an action-potential band, high-pass filtered at 300 Hz and sampled at 30 kHz, and a local field potential band, low-pass filtered at 1–250 Hz and sampled at 2.5 kHz. The angular position of the rotating stage was read from the encoder of the Newport controller and recorded, together with the synchronisation pulse generated by the IMEC probe, as digital lines on a National Instruments PXIe-6341 board sampled at 500 kHz. The sync signal was used to align the electrophysiological recordings to the stage rotation.

### Experimental protocol

The experimental protocol consisted of a sequence of seven 5-minute recording conditions, to assess neural activity under different vestibulo-visual contexts. For each condition, the motorized rotating stage played a sequence of clockwise and counterclockwise turning movements that covered 720°, with commanded peak angular velocities of 140°/second and peak accelerations of 200°/second² (Figure 1); the measured speed over the session had a median of 87°/s (99^th^ percentile 210°/s), robustly activating the vestibular system (Carcaud et al, 2017). The pseudorandom rotation pattern had been pre-recorded by manually turning the stage (similar to Preston-Ferrer et al. 2016), and aims to mimic naturalistic vestibular and proprioceptive stimulation (Angelaki and Cullen 2008). The same pre-recorded rotation sequence was replayed in every condition, so angular occupancy and angular-velocity statistics were identical across conditions. The stage movement was synchronized with electrophysiological data acquisition via rotary encoder signals A and B fed into SpikeGLX channels for subsequent analyses of stimulus-evoked neural responses.

**Figure 1.**
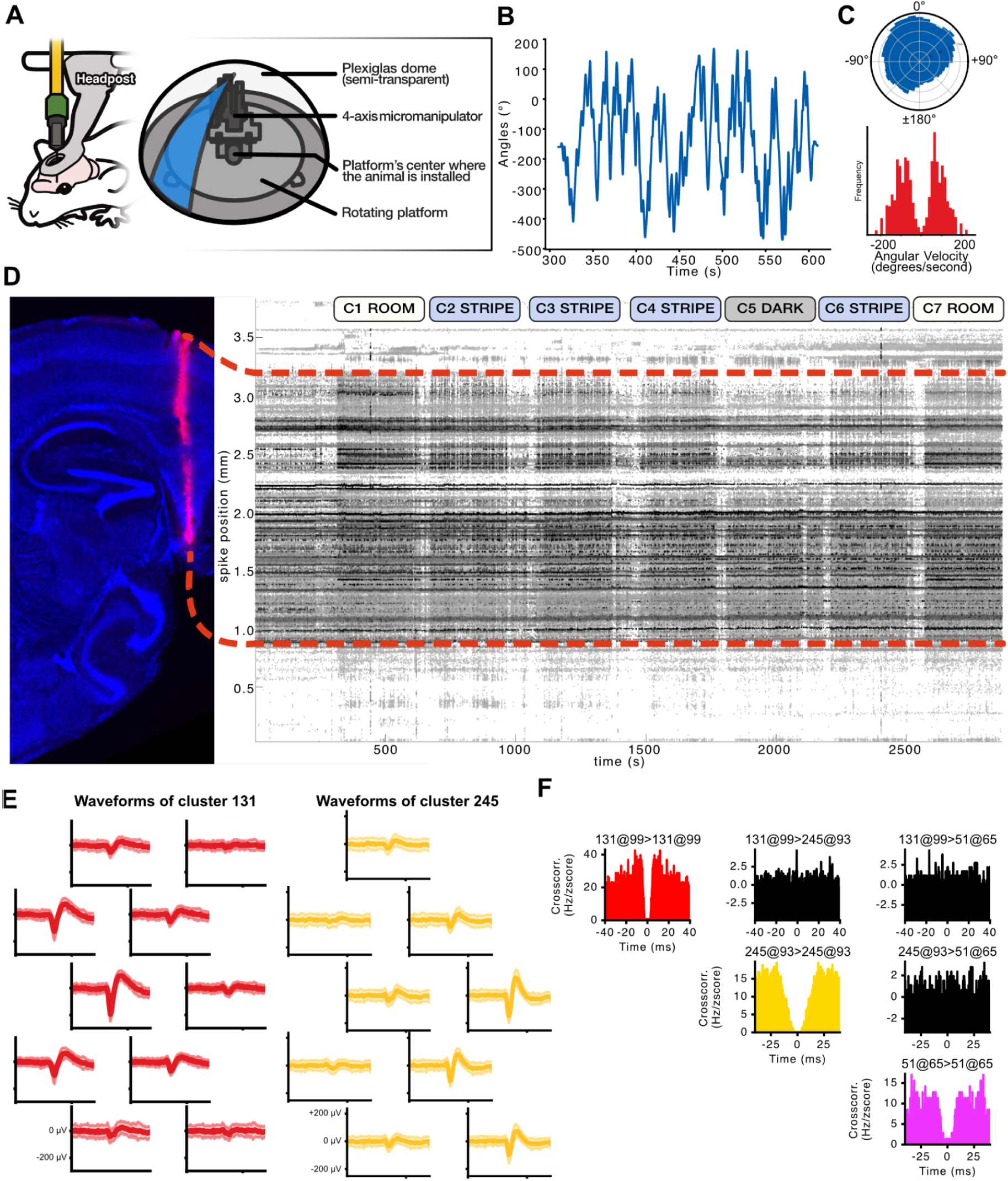
Experimental setup, anatomy, spike sorting. (A) Mice were head fixed at the center of the platform that was rotated around a central axis. A rotary encoder was used to read out the azimuthal angle of the platform. To control the visual surround, a dome with a visual cue (stripe) surrounded the platform in conditions C2, C3, C4 and C6. (B) Imposed rotation pattern for one 5-minute block (platform angle unwrapped), a pseudorandom sequence of clockwise and anticlockwise turns spanning ∼720°. (C) Statistics of the imposed rotation with angular coverage per 10° bin (top), and distribution of angular speed in deg/sec (bottom; median of 87°/s). (D) Left, typical probe track, labeled with DiI, crosses through the primary visual cortical layers 1 to 6 (VIS), and through the subicular complex (SUB), including pre-, post-, and/or parasubiculum and subiculum. Right, recording stability over time can be inferred from the drift map. Recording conditions C1 to C7 are indicated. (E) Spike waveforms appear on several contact sites, shown are units 131 and 245. (F) Auto-correlograms indicate well sorted units with refractory periods. Suppl. Fig 1-1, histologies for all animals #1 to 6 and their track reconstruction labels.

The sequence of conditions was as follows:

1. **Room Condition 1 (C1):** The initial 5-minute session was conducted in the cue-rich recording room environment, in dimmed light. The animal was exposed to the visual features of the recording room (corners, door, furniture).
2. **Stripe Condition 2 (C2):** The setup was covered with a dome-like screen surrounding the animal. A single blue luminous visual cue covering 30°. This cue was the sole visual reference during the rotation.
3. **Stripe Condition 3 (C3):** The visual cue was shifted by 90°, effectively relocating the reference point.
4. **Stripe Condition 4 (C4):** The visual cue was returned to its C2 position.
5. **Dark Condition 5 (C5):** No visual cue, to assess neural activity driven solely by vestibular inputs.
6. **Stripe Condition 6 (C6):** The visual cue was reintroduced at the same position as in C2 and C4, reinstating the visual reference point as before the dark condition.
7. **Room Condition 7 (C7):** The dome-like screen was removed, as in the initial room condition.

### Disorientation Periods

Between C2 and C3, C3 and C4, and C5 and C6, before changing the position of the stripe, the mouse underwent a 2-minutes disorientation period, where the motorized stage executed random bidirectional turning movements in the dark. This disorientation was implemented to prevent the mice from anticipating the subsequent cue position (Zugaro et al. 2001, 2003; Grieves et al. 2022). The 2-min duration is shorter than the 5-min and more used in Zugaro and colleagues; as such, any residual orientation across the gap may bias against the drift we observe between repeated cue presentations, making the SUB reinstatement residual (Figure 3B) a conservative estimate.

**Figure 2.**
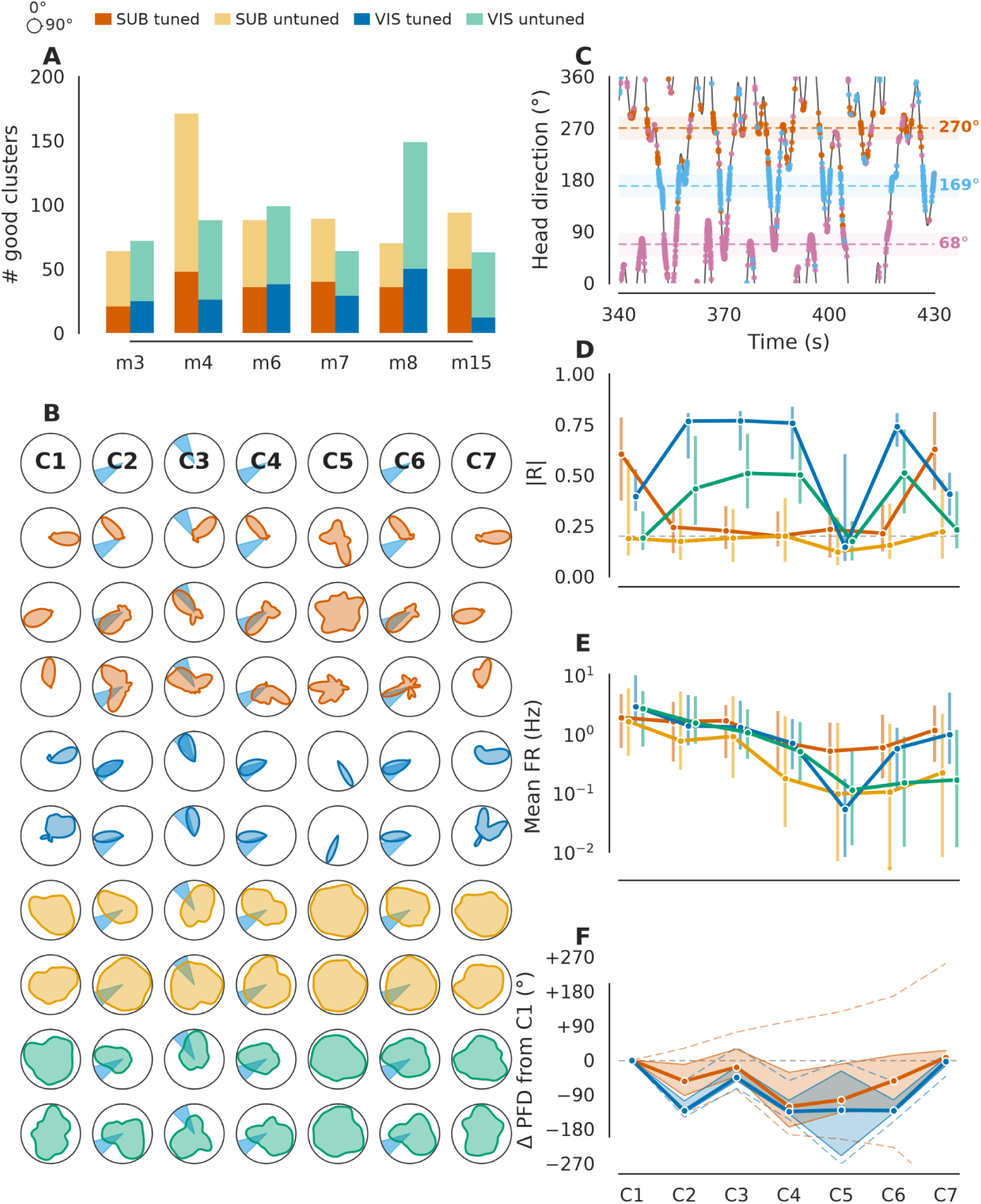
Directional tuning properties across conditions. (A) Histogram of SUB and VIS unit counts across six mice. Shown are all well identified units, grouped by region and by the selection criterion (tuned in C1 and C7): SUB directionally tuned, red; SUB untuned, yellow; VIS directionally tuned, blue; VIS untuned, green. (B) Polar plots of directional tuning for individual cells across seven conditions C1 to C7. The blue wedge indicates the position of the visual cue (present in C2-4, C6). The first three examples correspond to the cells shown in panel C. Same color code as in A. (C) Head direction trace (black) with the spikes of the three example cells overlaid; each dot is a spike, with one color per cell, and dashed lines mark each cell’s preferred firing direction. (D) Single cell tuning strength R_cell per group across conditions (group median across all identified units, with the dashed reference line making the selection threshold R_cell = 0.2). (E) Mean firing rates (Hz) for each group across conditions (log scale). (F) Change in preferred firing direction relative to each cell’s one C1 (ΔPFD in degrees), tuned groups only; group median over nested 10-90 % (shaded region) and 25-75 % (dashed line) quantile bands, the bands describe the spread of cells and not the uncertainty of the median. (B-F) is data from one example mouse (m4).

**Figure 3.**
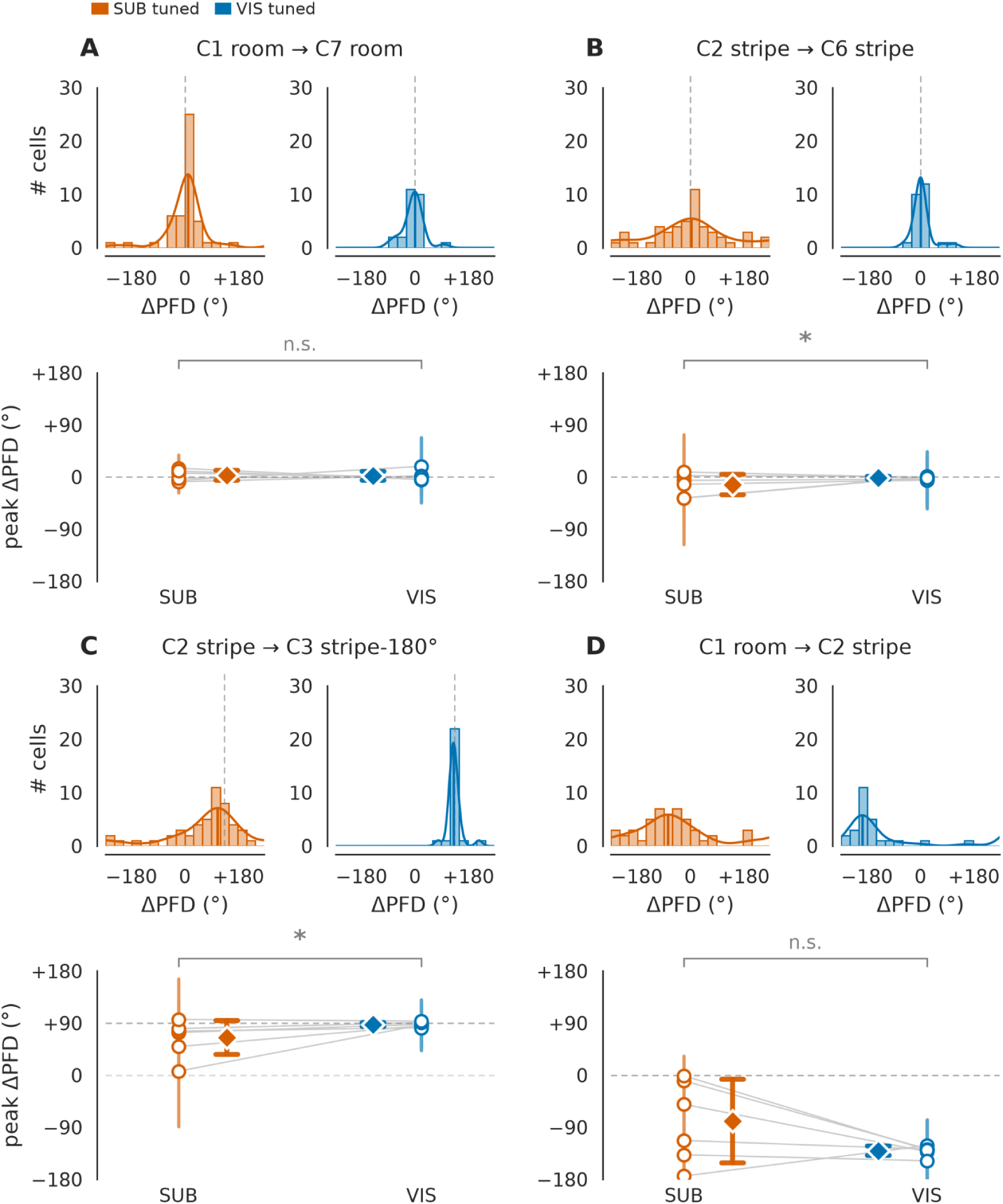
Directional tuning is stable across repeated exposures, and shifts systematically with cue manipulation. Four conditions pair, one panel each: (A) room reinstatement, between C1 and C7; (B) stripe reinstatement, between C2 and C6; (C) cue rotation of 90°, between C2 and C3; (D) room to stripe transition, between C1 and C2 (no expected shift defined). Per panel, top: distributions of per-cell ΔPFD for the example animal (m4), with SUB in red and VIS in blue, von mises fit (solid line) and the expected shift (dashed line). Bottom: aggregate across the six mice, each animal contributing the peak of its ΔPFD distribution (dot) with its 68% highest density interval (whisker), and the connecting lines pair each animal’s SUB and VIS peaks. Lane annotations give the concentration of the six per-animal peaks and the mean ± SEM absolute deviation from the expected shift.

### Data analysis

Electrophysiological data were pre-processed using CATGT for phase shift correction, noise reduction, artifact removal, and signal normalization to prepare the raw data for subsequent spike sorting. Spikes were sorted using Kilosort3 (Pachitariu et al. 2016, with the version-3 release), to detect and cluster extracellular action potentials across the 384 channels, then manually curated using Phy (Figure 1E-F).

### Linking Neural Activity to behavioural Data

The quadrature signals generated by the rotary encoder of the motorized stage were converted into angles, using a custom Python script. Spike times were temporally aligned with the corresponding stage angles, to correlate neural activity with the animal’s head direction. The HD trace was resampled onto a regular 100 Hz grid using nearest-neighbour interpolation. For 10.3% of grid samples, the nearest encoder event was more than 10 ms away (up to 122 ms in the worst case), limiting the effective temporal resolution of all direction-resolved analyses.

### Quantification of the angular tuning

Angular tuning of single neurons was quantified from the instantaneous firing rate and head direction, both sampled at 100 Hz (Taube 1995). Spike counts were computed in 10 ms bins and converted to firing rates (spikes/s). The HD tuning curve was then estimated at 1° resolution over the full circle using a circular Gaussian kernel (σ = 10°), such that observations at nearby directions contributed to the estimated rate according to their angular distance. Because the estimate is normalised by the summed kernel weight at each direction, it is occupancy-corrected by construction. Each angular tuning curve was then modelled using a von Mises distribution, with parameters corresponding to the preferred firing direction (PFD), concentration (κ), amplitude (peak firing rate), and baseline firing rate. Model parameters were optimized using bounded nonlinear least squares, and the proportion of variance accounted for (VAF) was used to evaluate the goodness of fit.

Normalized tuning amplitude (NTA) and the Rayleigh vector length of the tuning curve (R_cell) were used as measures of directional modulation and selectivity; when angular sampling deviated from uniformity, tuning curves were occupancy-corrected before R_cell was computed. Statistical significance of directional modulation was assessed using a non-parametric shuffle test. For each neuron, the spike train was circularly shifted relative to the HD trace to generate 1,000 surrogate datasets under the null hypothesis of no directional modulation. The Rayleigh vector length of the original data was then compared with the distribution of Rayleigh vector lengths obtained from the surrogate datasets, and neurons were considered significantly directionally tuned when the observed value exceeded the 95th percentile of the null distribution (p ≤ 0.05). Circular shifts were used rather than IID shuffles, so that each cell’s firing rate autocorrelation is preserved.

### Classification of Directionally Tuned Units and Preferred Direction of Firing

Units were classified as directionally tuned if they exhibited significant directional modulation (p ≤ 0.05) and R_cell ≥ 0.2 in both the first (C1) and last (C7) cue-rich room conditions. This intersection is substantially more restrictive than either endpoint alone (for example, in one animal 92 of 171 subicular units met the criterion in C1 and 68 in C7, but 48 in both) and ensures that units were detected and directionally tuned across the whole recording. The PFD was defined as the phase of the Rayleigh resultant vector of the tuning curve; it agrees closely with the fitted von Mises peak for well-tuned cells (median difference 5.2° in C1 or C7) and, unlike the fit, degrades for weakly tuned cells. Only units meeting the criterion in both C1 and C7 were included in subsequent analyses unless stated otherwise.

### Histology

After the recording session, animals were anesthetized by i.p. injection of a ketamine/xylazine mixture (100 mg/kg ketamine and 10 mg/kg xylazine). An intracardiac perfusion with 0.1 M PBS was followed by perfusion with 4% paraformaldehyde. Brains were then removed, stored overnight in paraformaldehyde at 4°C and sagittal sections were cut at 100 µm with a vibratome (LeicaVT1000S), washed in DAPI (15 minutes), then in PBS (two changes, 15 minutes each) before being mounted on coverslips using ProLong Gold Antifade (Invitrogen).

### Imaging and probe track reconstruction

Mounted brain sections were imaged using an epifluorescence microscope (Nikon) equipped with appropriate filters. The acquired images were aligned and matched to the reference mouse brain atlas provided by the Allen Brain Institute using the SHARP-Track tool (https://github.com/cortex-lab/allenCCF). Serial sections were compiled to reconstruct the three dimensional trajectory of each Neuropixels probe. The probe track was reconstructed so that recording sites could be matched to anatomically defined regions, for each mouse. Only brains with clearly identifiable probe tracks corresponding to the intended recording regions were included in the final analysis (Figure 1D, supplementary Figure 1.1). Units were assigned to VIS when the reconstructed recording site fell within the primary visual cortex (VISp, any layer). Subicular complex units (SUB) had their recording sites in either pre-, post-, parasubiculum or subiculum (Supp. Fig. 2-2).

### Shift distributions and reinstatement

For each condition pair, per-cell ΔPFD was computed as the circular difference of PFDs. Per animal, the mode of the ΔPFD distribution was estimated with a von Mises kernel density (histogram fallback for small samples) and reported with its 68% highest-density interval. Re-alignment was quantified as the fraction of cells whose ΔPFD lay within ±20° of the expected shift, computed both over all selected cells and restricted to cells significantly tuned in both conditions of the pair. Coherence of the shift was quantified as R_shift, the resultant Rayleigh vector length of the ΔPFD distribution, which is defined for all pairs between two conditions including those without an expected shift.

### Population tuning and stripe preference

Population firing rate maps were created for each condition. For each condition, maps were generated by stacking all significantly tuned neurons’ normalized head direction (HD) tuning curves. Each curve was normalized to its own firing rate range and aligned from -180° to +180°, so they could be plotted and sorted based on each neuron’s preferred firing direction (PFD) in the first light condition.

Population polar plots show the mean normalized tuning curve across cells with a null circular preference envelope: each cell’s tuning curve was rotated by an independent random offset and the population mean recomputed (1000 iterations), preserving every cell’s tuning depth and shape while destroying only the alignment between cells. The plotted envelope is the 5th to 95th percentiles of the population mean obtained across these iterations.. The population concentration (R_pop) was defined as the resultant length of the cells’ PFD unit vectors:

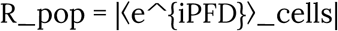

Since R_pop is positively biased for finite sample sizes, with an expected magnitude of approximately 1/√N under a uniform distribution, we estimated the chance level separately for each animal and condition by simulating 10,000 populations of *N* uniformly distributed PFDs, where N was that animal’s recorded cell count. The mean simulated resultant length was reported alongside the observed R_pop.

The stripe preference index (SPI) compared the arithmetic mean of each cell’s normalized firing rate across angular bins within a ±40° wedge centered on the stripe with that within the diametrically opposite (“shadow”) wedge.:

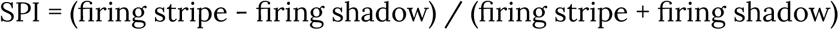

which ranged from -1 (opposite preference) to +1 (stripe preference). SPI is undefined in darkness (no landmark). The angular position of the door in the room conditions was taken as a single calibrated value (20°) for all sessions; cue-relative quantities in C1 and C7 are therefore approximate, whereas stripe positions in C2-C4 and C6 were set by the projector. SPI values are reported as one median per animal and compared between SUB and VIS with a paired Wilcoxon signed-rank test across the six mice.

### Population coherence

Shared temporal structure within each condition was quantified by calculating Pearson correlations between the 100-ms-binned firing-rate traces of all pairs of cells over a fixed 300-s window starting at the onset of each condition. Cells were included only if they fired at least 30 spikes in every condition, ensuring that the same cells and cell pairs were included and identically ordered in the correlation matrices across conditions. Pairs were classified as similarly tuned (“aligned”, ≤20° PFD difference) or dissimilarly tuned (>20° PFD difference) based on their PFD; unless stated otherwise, the pair labels derive from each condition’s own PFDs in Figure 5 and from the C1 PFDs, carried unchanged into every condition, in Supp. Fig. 5-1. The strength of correlated activity in aligned versus non-aligned pairs was compared within animals by permutation and across animals by Wilcoxon signed-rank on per-animal means. Similarity of correlation structure across conditions was quantified by calculating the Pearson correlation between the strict upper triangles of the corresponding cell–cell correlation matrices. Split-half reliability of each condition’s matrix (interleaved bins, Spearman–Brown corrected) and attenuation-corrected similarities are reported in Supp. Fig. 5-2.

**Figure 4.**
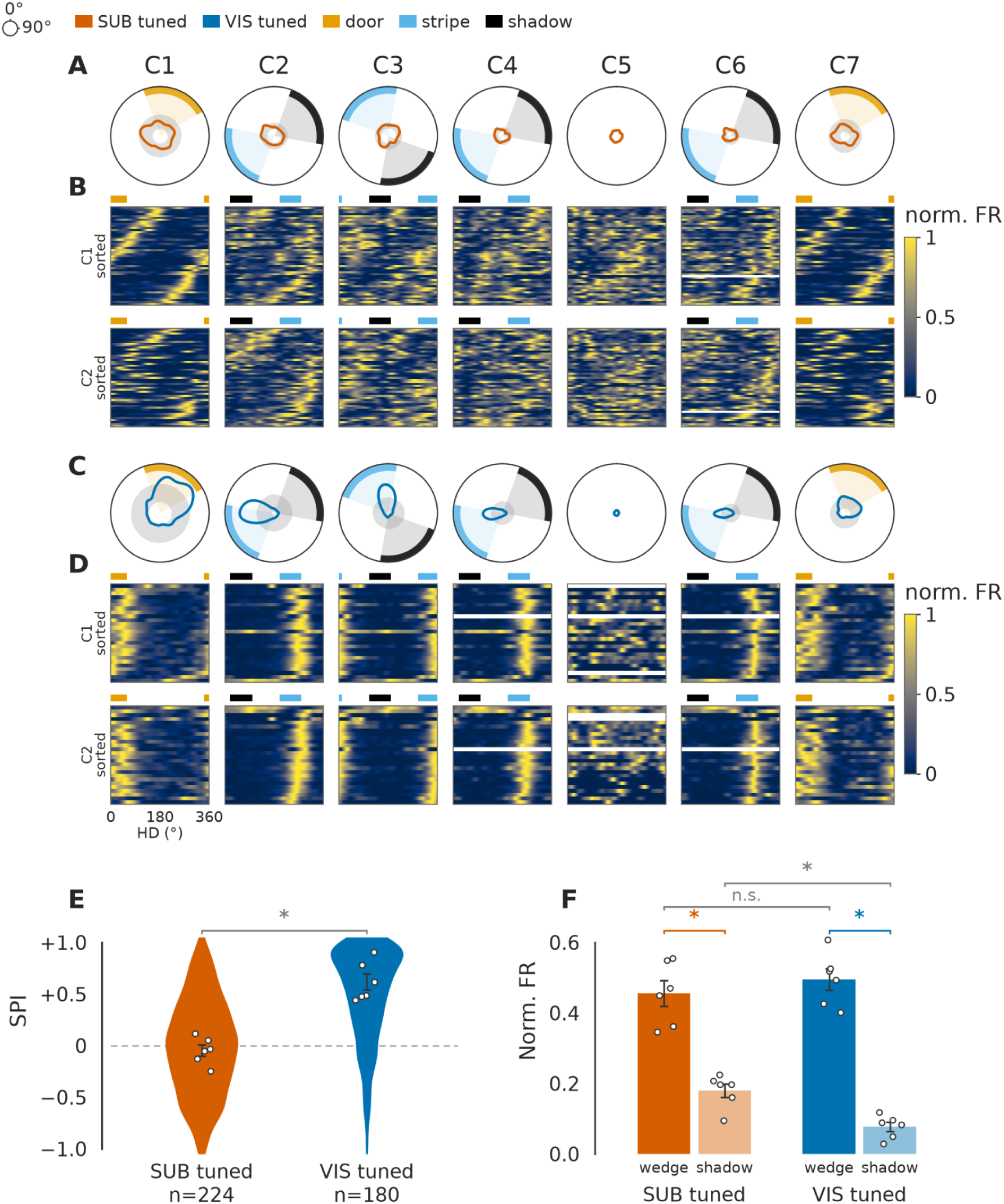
Uniform distribution of preferred directions in subicular complex cells and cue alignment in visual cortical cells. (A) Ordered head-direction tuning curves of the tuned SUB cell population (each row corresponds to a cell) from one example animal (m4) and across conditions. Top: cells ordered by the preferred head direction in the room (C1). Bottom: cells ordered by preferred head direction in stripe (C2); the cell order is held across all seven columns. Amber yellow bands indicate the door position in the room (C1, C7), blue bands indicate the stripe position, and black bands indicate the opposite position (“shadow”). (B) Polar plots of the mean activity of tuned SUB cells across conditions. Doughnut-shaped grey areas indicate the circular shift null envelope (each cell’s curve rotated by an independent random offset, 1000 iterations, 5th–95th percentile interval), with a mean curve inside the envelope indicating preferred directions no more aligned than chance; indicated are the positions of the door (amber), stripe (blue), and the side opposite to the stripe (“shadow”, black). (C) Same analysis as in panel A for tuned VIS cells. (D) Same analysis as in panel B for the tuned VIS cells. (E) Stripe preference index distributions (SPI; violin plots) for tuned SUB (red) and tuned VIS (blue) cells. Positive values indicate preference toward the stripe; negative values indicate preference toward the shadow. SPI is undefined in darkness. (F) Mean normalized firing rates (mean ± SEM) at the stripe wedge vs. the shadow positions. The SUB cells tuned to the stripe are shown in red, and the VIS cells tuned to the stripe in blue; shadow responses are shown in lighter red and lighter blue. All statistics are paired Wilcoxon signed-rank tests across the n = 6 mice.

**Figure 5.**
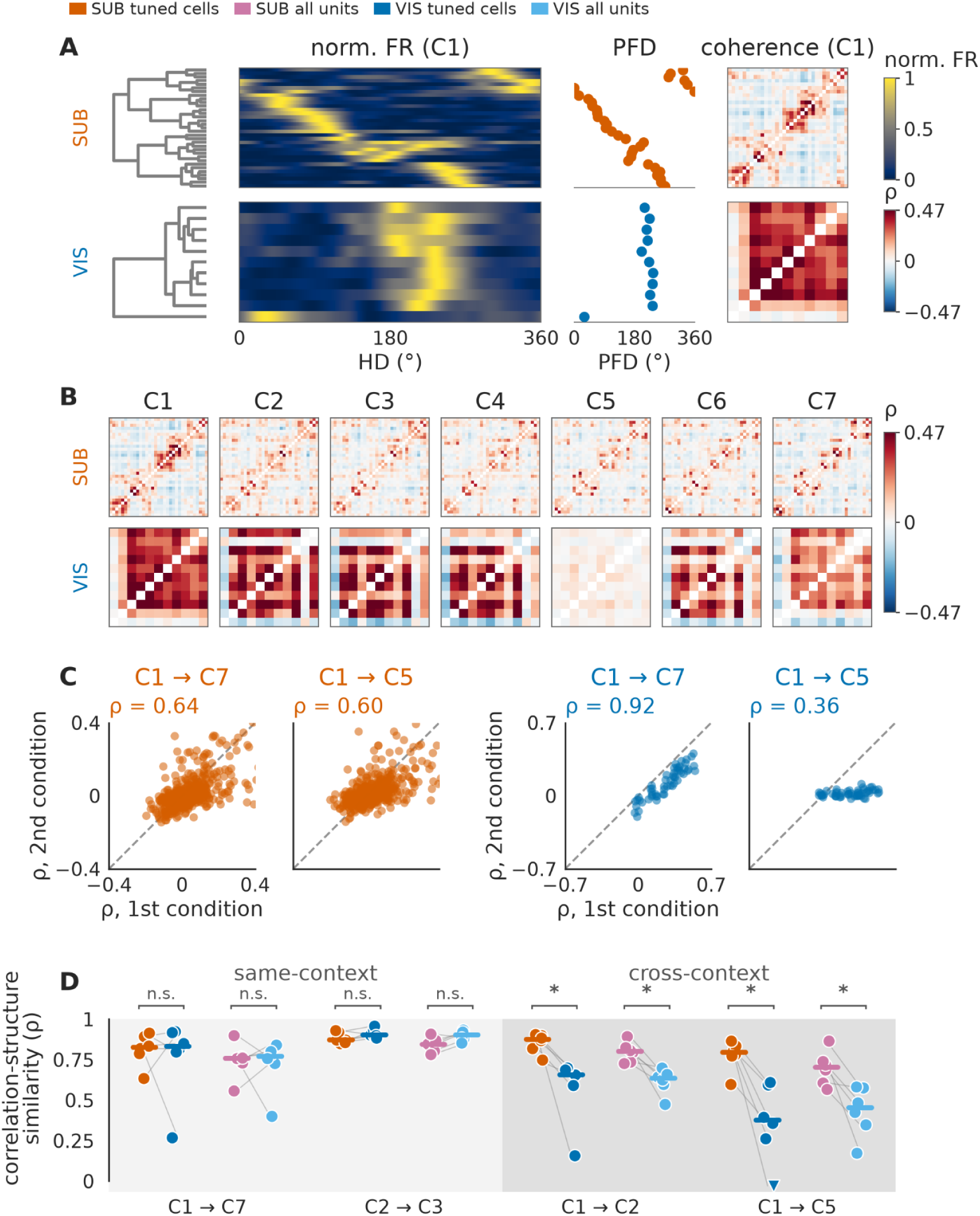
Network structure and coherence across conditions. (A) Clustering and organisation of tuned units in one example animal (m4). From left to right: the hierarchical clustering dendrogram of SUB-tuned cells using Ward’s method on the z-scored C1 tuning curves; the normalized firing rate by angle for each cell; the preferred firing direction per cell and the signed pairwise correlation matrix (100-ms binned rates, 300-s window) between cells, using the same ordering. Bottom: same for VIS-tuned cells. Cells are included only if active in every condition (termed : complete cases), so counts differ slightly from Figure 2. (B) Pairwise correlation matrices across all conditions using the ordering derived for C1 (see A). Top: SUB-tuned cells; bottom: VIS-tuned cells. (C) Scatter plots of cell-pair correlation values between two conditions, for one example animal (m4). SUB-tuned pairs, red; VIS-tuned pairs, blue. (D) Distribution of Pearson’s ρ values across condition pairs, for n=6 mice (one dot is one animal). Plots show SUB-tuned (red), VIS-tuned (blue), all SUB units (pink), and all VIS units (light blue), highlighting coherence differences within and across contexts. Light gray block, coherence for pairs in same-context, dark gray block, cross-context comparison. Paired Wilcoxon signed-rank across mice.

### Population structure and decoding

Head Direction and angular velocity was decoded from the population activity of neurons either from SUB, or, from VIS, or from both. Spikes were filtered with a gaussian low-pass filter (σ = 200 ms). The nonlinear decoder was trained with ε-support-vector regression (SVR) with radial basis function (RBF) kernel on a contiguous 225 seconds of trajectory, operating directly on the smoothed spike space. Testing was done on the remaining 75 s. The 225-s training window was placed at seven different positions within the block, with a 3-s guard band separating training and test data. Predictions from the resulting overlapping test windows were pooled to obtain a single decoding estimate and were not treated as independent observations. For head direction, we separately trained to predict sine and cosine of head direction (necessary because of the circular nature of this variable), then combined results using arctangent. For angular velocity, the decoder was trained to predict velocity and z-scored on training statistics (SVR hyperparameters are absolute-scale).

Signed decoding error as a function of true head direction was computed by smoothing the decoded direction as a circular mean weighted by proximity in true direction (circular Gaussian, σ = 10°) and taking its signed circular difference from the true direction. Stripe blocks were rotated by their own landmark angle onto the common 90° stripe position (240° in room coordinates) before pooling; room blocks were not rotated. Modulation was quantified per animal as the peak-to-trough amplitude of the smoothed curve and as the amplitude and phase of its first circular harmonic, each tested against a null of 1000 circular shifts of the signed-error series relative to the head-direction series within held-out windows, preserving the autocorrelation of both series. Angular occupancy of the cue and shadow wedges is reported alongside. To compare landmark placements, the same quantities were also computed per condition without rotation, and the anchor of the deformation (the descending zero of the first harmonic) was expressed relative to the cue.

For cross-condition generalisation (Supp. Fig. 7-1), one model per animal, region and training condition was fitted on the whole training block and applied unchanged to every other block; the constant offset between decoded and true direction was estimated as a circular mean, and error was re-measured after removing it.

**Figure 6.**
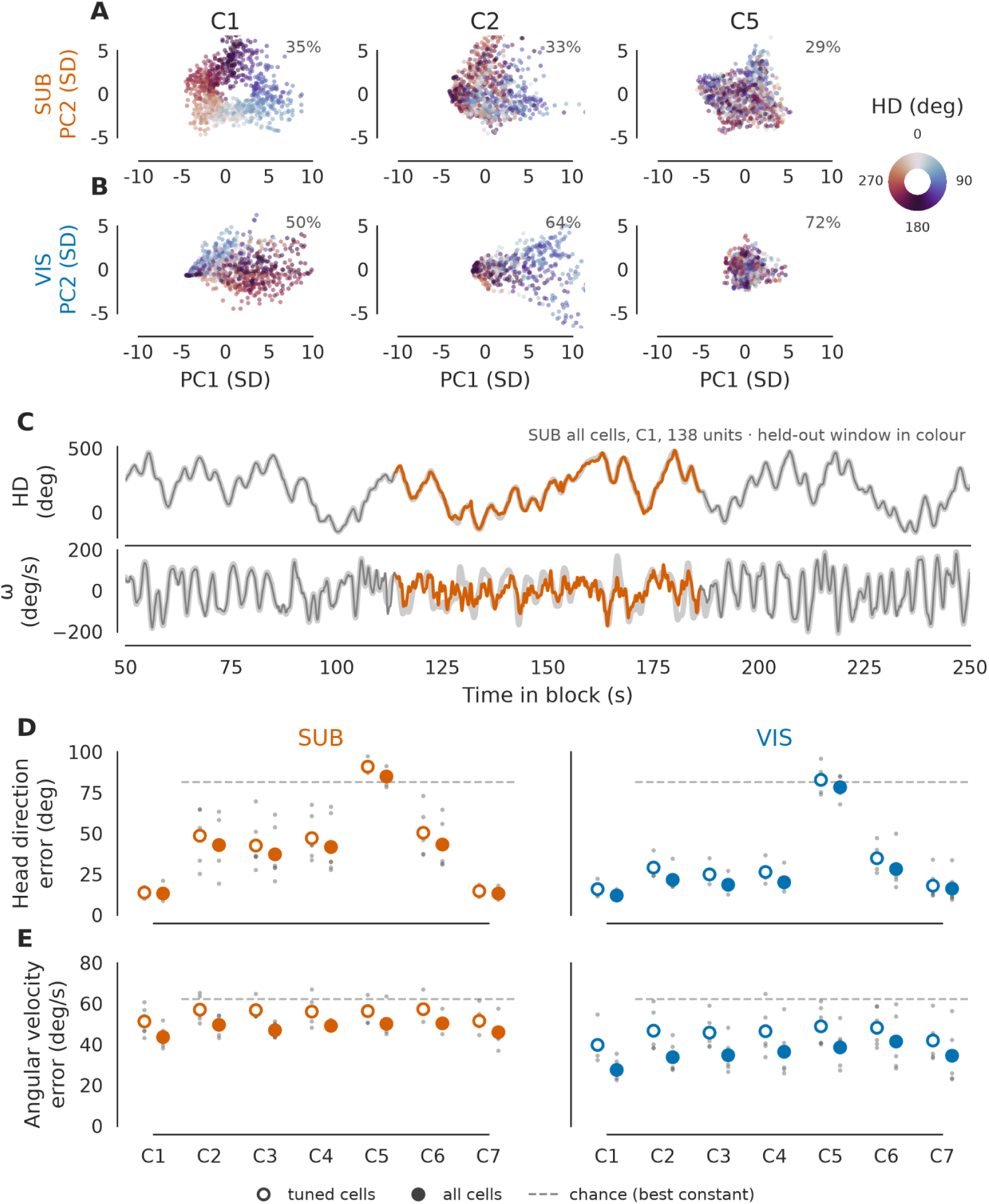
Population geometry and nonlinear decoding of head direction and angular velocity. (A) PCA projections of smoothed SUB population activity in Room (C1), Stripe (C2) and Darkness (C5), each dot colour-coded by the animal’s true head direction (example animal m4). The first two PCA components are shown. PCA is used for visualisation only; decoding operates on the full population activity. (B) Same as (A) for VIS. (C) The decoder on one example block (SUB, all cells, C1, example animal m4): decoded head direction and angular velocity (colour) against the true trajectory (black); held-out test window in colour, training window in grey. (D) Median circular absolute error of head-direction decoding for C1-C7, for SUB and VIS, with the decoder trained on all recorded cells (filled) in the respective condition or on tuned cells only (open); one point per animal, bar = mean across the six mice, dashed line = the measured chance floor (best constant prediction, 81.4°, identical across conditions because the rotation sequence is replayed). (E) Same as (D) for angular velocity (floor 62.1 °/s). Statistics: paired Wilcoxon signed-rank across mice. Decoding the combined SUB+VIS population improved on either region alone only through its larger cell count: subsampled to a single region’s count, the joint population never outperformed the better single region (Methods; Results).

**Figure 7.**
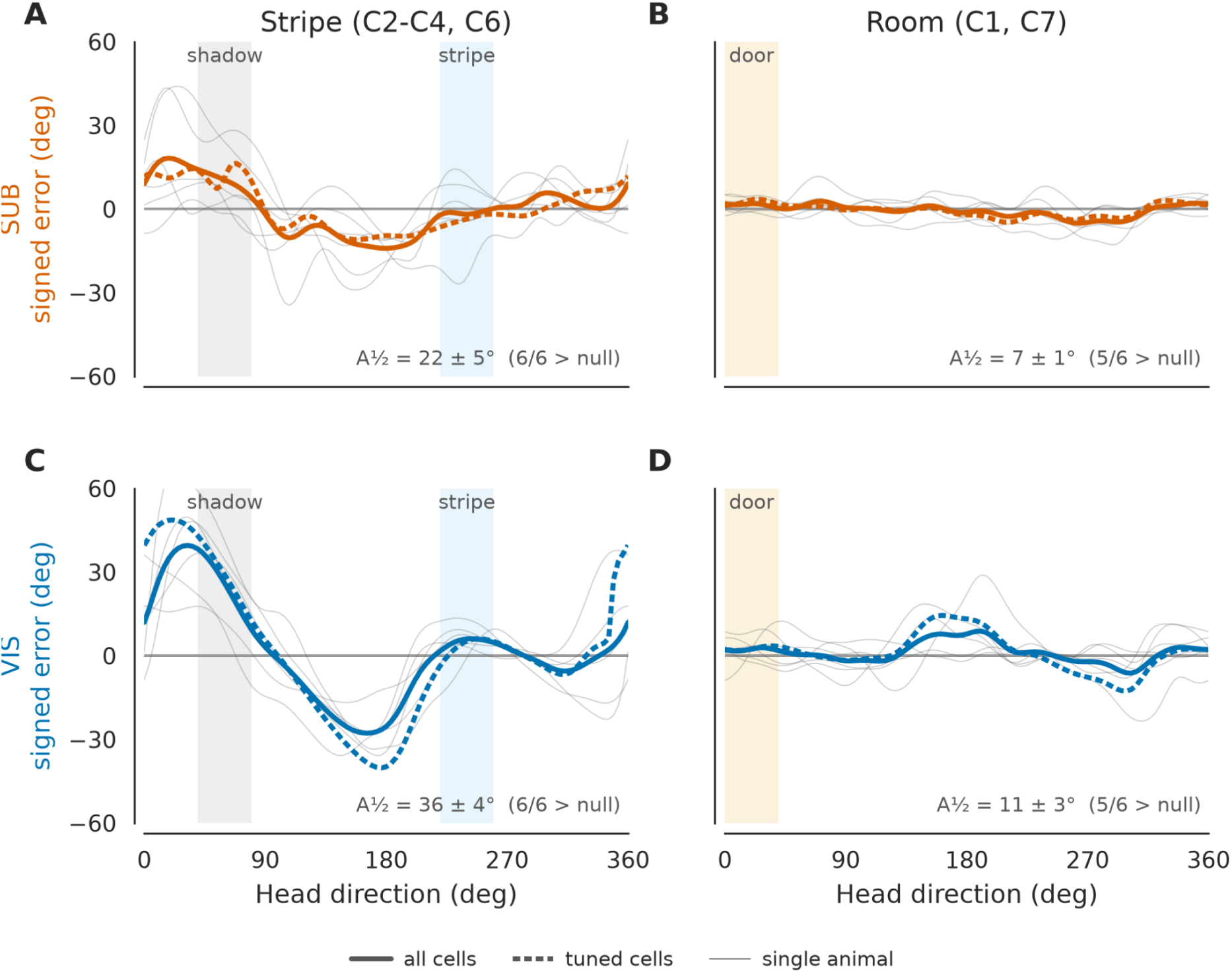
Accuracy of decoding as a function of stripe direction. (A) Median signed decoding error (°) as a function of Δ to stripe (°), for SUB tuned cells. Shaded bands indicate the stripe ± 20° region (light blue) and the shadow position (opposite direction, gray shaded region). Within-condition decoding, including stripe C2-C4 and C6, aligned by stripe position. (B) Same analysis for SUB tuned cells in room (C1, C7). The shaded band indicates the door position. (C) Same analyses as in panel A for VIS tuned cells in stripe (C2-C4, C6). (D) Same analyses as in panel B for VIS tuned cells in room (C1, C7). Data from SUB populations are shown in red, and VIS populations in blue. Gray lines are individual mice.

### Statistics

All comparisons across animals used two-sided Wilcoxon signed-rank tests over n = 6 mice, reported with the test statistic W and the matched-pairs rank-biserial correlation r. Quantities were collapsed to one value per animal before testing, and cells were never treated as independent samples. All within-animal tests were circular shift permutation tests (≥1000 shifts, seeded), which preserve temporal structure and autocorrelation of the spike train; independent shuffles were never used. With n = 6 the smallest attainable two-sided signed-rank p is 0.031, so p = 0.031 with W = 0 indicates a unanimous effect across all six animals. Because familywise correction over three or more such tests cannot reach significance regardless of effect size, one primary contrast was pre-specified per supplementary analysis and all remaining tests are reported uncorrected and labelled exploratory.

## Results

We investigated directional signals in the subicular complex (SUB), comprising pre-, post-, parasubiculum, and subiculum proper, and the primary visual cortex (VIS) in head-fixed mice (Figure 1A-D). Animals were passively rotated across seven conditions, including cue rich room conditions (C1 & C7), conditions with a single visual cue (C2-4, C6), and a darkness condition (C5). Across six animals, 576 well isolated units were obtained from SUB and 535 from VIS (Figure 2A). The imposed rotation sampled all azimuthal directions near-uniformly in every condition (Figure 1C), so no comparison below is confounded by directional sampling. The results below examine classical criteria that define head direction cells, including directional tuning, stability over time, and cue control.

### Directional tuning across regions and sensory contexts

To link neural data to head direction, spike trains were aligned to the instantaneous head direction angle of the animal and the directional selectivity was quantified by the Rayleigh vector length of the tuning curve (R_cell). In the cue-rich room conditions C1 and C7, many neurons in both SUB and VIS displayed significant directional modulation (R_cell ⩾ 0.2, p < 0.05) with well defined tuning curves evident in both individual cell and population average tuning (Figure 2A-C). The percentage of directionally tuned neurons was 53.6 ± 4.0 % in SUB and 54.1 ± 3.7 % in VIS in C1 (mean ± SEM across n=6 mice, fraction of well-isolated units of each structure). Preferred tuning was stable over time: 77 ± 5 % of tuned SUB neurons and 69 ± 10 % of VIS neurons maintained their preferred firing direction from first (C1) to last room condition (C7) within ± 20° of their original preferred firing direction (Supp. Fig. 3-1). For subsequent analysis we selected a subset of units in SUB (n = 231) and in VIS (n = 180) that satisfied the criteria of directional tuning in both room conditions C1 and C7 (|R| ⩾ 0.2 and p < 0.05 in each), ensuring that the considered units were active and directionally tuned at the beginning and at the end of the recording period.

### Reinstatement across repeated cue configurations

Re-exposure to a familiar environment during the protocol let us examine the stability of directional tuning over time and whether the original HD alignment was retained. Between the two room exposures (C1 vs. C7), both regions returned to their original directional maps with similar fidelity. The per-animal peak of the ΔPFD distribution deviated from zero by 8.1 ± 1.8° in SUB and 5.0 ± 2.8° in VIS (n.s. different; Wilcoxon signed-rank, n = 6, W = 6, p = 0.44), with population concentrations restored to their C1 values (C7: SUB R_pop 0.198 ± 0.036 against simulated chance floor of 0.148; VIS 0.571 ± 0.065 against 0.174; Figure 3A). The stripe reinstatement (C2 vs. C6) dissociated the regions, with VIS re-aligning to the reinstated stripe (peak deviation 1.9 ± 0.9°) whereas SUB carried a residual mis-alignment of 17.0 ± 6.3° with a systematic bias (circular mean of -13°), larger than the VIS residual in every animal (W = 0, p = 0.031, r = 0.88; Figure 3B). Since the stripe returns to its exact previous position, the visual re-alignment is expected from the stimulus geometry alone; as such, the subicular residual instead measures how far the internal directional estimate drifted during the dark and short disorientation periods. Both regions seem to maintain the association between landmark and directional map, but the subicular map is additionally updated by an internal estimate that accumulates error when unanchored (Valerio and Taube 2012; Peyrache et al. 2015).

### Cue anchoring and rotation of preferred direction

To assess how a single visual cue controls tuning, we compared PFDs before and after shifting the stripe by 90° (C2 vs C3) and after returning it to its initial position (C2 vs C4; Figure 2B, F; Figure 3C). Following the 90° shift, the visual cortical population rotated with the cue. The per-animal peak of the ΔPFD distribution was of +87.7° for a commanded +90°, deviating by only 3.7 ± 1.0°. The subicular population also followed the cue but under-rotated in some animals (peak +65.2°; deviation 28.3 ± 12.0°; SUB vs. VIS deviation: W = 0, n = 6, p = 0.031, r = 0.88). The shift distributions per animal were coherent in both regions (concentration of the six per-animal peaks: SUB 0.88, VIS 1.00), so the subicular under-rotation is a property of the population; in other words, the population moved together and landed short of the cue-predicted direction. Under passive rotation, the vestibular record of the platform’s actual motion is highly reliable, and thus the observed undershoot may occur as the subicular estimate is a compromise between the rotated visual frame and the un-rotated vestibular one (J. Goodridge and Taube 1997; Knierim et al. 1998; Zugaro et al. 2003). Following the backshift (C2 vs C4), 76 ± 6 % of the tuned VIS cells and 34 ± 6 % of SUB cells returned to within ±20° of their original preferred direction (95 ± 2 % and 51 ± 9 % when restricted to cells significantly tuned in both conditions; SUB below VIS in every animal, W = 0, p = 0.031). Note that only 22 to 31 % of subicular good units are significantly tuned under the single stripe (against 54 to 70 % in VIS; Supp. Fig. 2-1), so unrestricted re-alignment fractions score many subicular cells on PFDs estimated from weakly modulated curves, and the regional contrast survives the restriction at roughly half its magnitude (Supp. Fig. 3-1).

### Context-dependent shifts and tuning stability

Transitions between contexts produced region-specific reorganisation. Moving from the room to the dome (C1 → C2; Figure 2B, F and Figure 3D), the visual cortical population shifted consistently across animals, by -131 ± 4°. This shift was close to the -130° predicted from the change in landmark angle between the door in the room and the stripe in the dome. This correspondence is consistent with the hypothesis that visual cortical representations align to salient landmarks in both environments. The subicular population did not shift coherently across animals. Its per-animal shift peaks spanned nearly the full circle (concentration of the six peaks 0.45). The room and the stripe are therefore interchangeable reference frames for VIS, related by a simple rotation, while for SUB they are distinct contexts between which the re-mapping is established animal by animal. Tuning quality moved in opposite directions across the same transition, with the subicular tuning degraded (median R_cell 0.73 ± 0.04 in C1 to 0.36 ± 0.05 in C2; tuning width 65 ± 5° to 79 ± 3°; Fig. 2D) whereas the visual cortical tuning sharpened (R_cell 0.41 ± 0.02 to 0.55 ± 0.06; width 80 ± 2° to 69 ± 3°; Fig. 2D). Consequently, tuning width was narrower in SUB than in VIS in the room (W = 0, p = 0.031, r = 1.00) but the ordering reversed in every stripe condition. Together, these findings suggest that a single high-contrast stripe sweeping across the visual field may provide a particularly effective stimulus for eliciting directional responses in the visual cortex, whereas the subicular directional code may be most precisely anchored when multiple stable landmarks are available.

### Firing rate modulation across conditions

The firing rate modulation across conditions also followed the same dissociation between the two regions (Figure 2E and Supp. Fig 2-1). Peak firing rates of tuned SUB cells decreased from 8.3 ± 1.3 spk/s in the room (C1) to 3.6 ± 0.6 spk/s under the dome (C2), significant in all six animals (Wilcoxon signed-rank, n = 6, W = 0, p = 0.031, r = 1.00), and to 2.4 ± 0.4 Hz in darkness (C5). In contrast, VIS firing rates were consistently higher and showed little change from the room to the stripe condition (C1, 11.9 ± 1.2 to C2, 10.1 ± 1.3 spk/s), but decreased markedly in darkness (2.1 ± 0.5 spk/s). Under a single stripe, therefore, SUB cells showed a concomitant reduction in firing rate and directional tuning, whereas visual cortical cells maintained relatively stable firing rates while retaining directional tuning. Removing all visual inputs (C5) collapsed VIS firing rates. Rather than two gain-control mechanisms operating on a common directional signal, the pattern indicates two different signals: one landmark-dependent internal estimate in SUB, and a second stimulus-driven response in VIS for which the single stripe is not a degraded room but an optimal stimulus.

### Directional tuning in darkness

Passive rotation in darkness (C5) collapsed single cell directional tuning in both regions (Figure 2D and supp. Fig. 2-1). The fraction of directionally modulated cells fell sharply relative to the room (SUB: 3.4 ± 0.8 % vs 53.6 ± 4.0 %; VIS: 2.7 ± 0.9 % vs 54.1 ± 3.7 %; Wilcoxon signed-rank, n = 6, W = 0, p = 0.031 for each region), and single cell tuning strength approached its floor (R_cell: SUB 0.162 ± 0.015; VIS 0.138 ± 0.016), with preferred directions widely redistributed (Figure 2F).

At the population level, however, the two regions diverged in opposite directions. Continuous attractor models predict that in the absence of landmarks an internally generated HD signal persists but drifts through angular space. Consistent with this prediction, the concentration of preferred directions across cells fell in VIS (R_pop 0.563 ± 0.052 in C1 to 0.219 ± 0.051 in C5; W = 0, p = 0.031) but rose in SUB (0.179 ± 0.052 to 0.436 ± 0.092; W = 0, p = 0.031), and the previously tuned subicular ensembles concentrated toward a direction that differed in every animal and bore no relation to the room frame, as the residual alignment exceeded its circular shift null in four of six mice for SUB and two of six for VIS (Supp. Fig. 4-1). This is the population-level signature of an intact but un-anchored attractor as, with the landmark removed, the activity bump is free to drift and dwell, and whatever direction it occupies is imprinted coherently on every cell’s tuning curve (Skaggs et al. 1995; Zhang 1996; Ajabi et al. 2023).

However, two caveats need to be considered. First, in darkness most cells are weakly modulated and each PFD is estimated from a nearly flat curve, so slow drift within the block will itself inflate the apparent concentration. It is why we report the curve-based circular shift test rather than a point-based Rayleigh test. Second, the concentration says nothing about a fixed environmental orientation, as the compass continues to run but no longer points anywhere in particular.

### Population level analysis of preferred firing directions

We next asked, are the preferred firing directions of tuned SUB and VIS cells distributed uniformly across 360°? Ordering firing rate matrices by each neuron’s preferred direction in the room (C1) produced a diagonal that was preserved in C7, degraded under the stripe conditions and in darkness; ordering by the C2 PFD restored the diagonal alignment in the stripe conditions (Figure 4A). In the room, the subicular population showed no PFD anisotropy beyond chance, with R_pop 0.179 ± 0.052 in C1 and 0.198 ± 0.036 in C7 against a simulated chance floor of 0.148 (only one of six animals exceeded its own floor in either condition), in other words the population mean tuning curve remained inside the circular shift null envelope in the room conditions (Figure 4A). Under the single stripe, departures of the subicular population from the envelope were weak and inconsistent (up to three of six animals per stripe condition; R_pop 0.217–0.265; Supp. Fig. 4-1). Visual cortical population activity, on the other hand, was strongly and consistently concentrated in all light-on condition in the direction of the salient visual cue, that is toward the door in the room (R_pop 0.563 ± 0.052 in C1, all six animals above their floors) and toward the stripe under the dome (0.757 ± 0.060 in C2), with reduced activity in all other directions (Figure 4C-D). The population peak was displaced away from the stripe center (see Figure 7 for the corresponding displacement of the decoding error). These results thus show a near-uniform directional tiling of the environment in SUB that is weakly remapped by a single salient cue, against a landmark-locked distribution in VIS.

To quantify the influence of the presence of a single salient landmark on the distribution of preferred firing directions, we calculated a stripe preference index (SPI). This measure compared the normalized responses in a ±40° wedge around the stripe (total width : 30°) center to those in the opposite direction, the “shadow”. The SPI for SUB was close to zero (median SPI = -0.041 ± 0.058, for all stripe conditions, n = 6 mice), as expected for its homogeneous distribution of preferred firing directions across 360°. The VIS distributions were shifted to positive values, indicating a preferred activation around the stripe (median SPI = 0.619 ± 0.076; SUB vs. VIS Wilcoxon signed-rank, n = 6, W = 0, p = 0.031, r = 1.00; for all stripe conditions, n = 6 mice, Figure 4E).

Analysis of the normalized firing rates at the stripe and the shadow directions sharpened this picture (Figure 4F). Among cells firing preferentially toward the stripe (SPI > 0), both regions responded more toward the cue than toward the shadow (SUB 0.458 ± 0.036 vs 0.180 ± 0.019; VIS 0.496 ± 0.030 vs 0.079 ± 0.013; W = 0, p = 0.031, r = 1.00 within each region). The single cue modulates the subicular population too, by a factor of 2.5. The regional difference lies not at the cue, where the two populations were statistically indistinguishable (W = 9, p = 0.84), but at the shadow (W = 0, p = 0.031), where subicular cells retained 39 % of their cue-directed response and visual cortical cells only 16 %. A stimulus-driven interpretation predicts the visual cortical asymmetry directly, since nothing is present at the shadow to drive a visual response. The substantial subicular shadow response is instead what the rotational symmetry of ring attractor connectivity predicts (K. Zhang 1996; Knierim and Zhang 2012).

### Pairwise correlations and population coherence

Pairwise co-fluctuations of neuronal firing provides a measure of functional coupling that would be expected for an internally organized attractor network structure. Here, we sorted tuned units by hierarchical clustering of their C1 tuning curves and computed signed correlation matrices of 100-ms binned activity for each condition, keeping the same cell pairs and ordering across all conditions (Figure 5A-B). In the room (C1), both regions showed a clear dependence of pairwise correlations on tuning similarity: pairs with similar PFDs (ΔPFD ≤ 20°) exhibited positive co-fluctuations, whereas correlations between dissimilarly tuned pairs were around zero (SUB aligned vs non-aligned: 0.211 ± 0.025 vs 0.010 ± 0.011; VIS: 0.219 ± 0.026 vs 0.062 ± 0.023; within animal circular shift permutation p < 0.01 in all six animals; across animals W = 0, p = 0.031 for each region). Expressed as a continuous function of pairwise tuning difference, correlation decreased from approximately +0.21 for ΔPFD < 20° and progressively approached zero as tuning differences increased, similarly in both regions (Supp. Fig. 5-1).

This tuning-ordered coupling structure dissociated the regions once the scene changed. Pair labels in the following derive from each cell’s C1 preferred direction, carried unchanged into every condition, so the statement is that coupling ordered by the lit-room tuning survives the manipulation. This fits the specific prediction of an attractor whose connectivity defines the code, and not its anchoring (Skaggs et al. 1995; Zhang 1996; Chaudhuri et al. 2019). Under the single stripe, the subicular alignment contrast persisted at every condition (aligned minus non-aligned Δρ = 0.11–0.12 at C2, C3, C4 and C6; W = 0, p = 0.031 at each). In darkness, the subicular gradient retained 75 % of its room span (aligned pairs 0.211 → 0.140; alignment contrast 0.201 → 0.111; aligned vs non-aligned at C5: W = 0, p = 0.031). On the other hand, the visual cortical gradient collapsed to 15 % of its span, with aligned VIS pairs falling from 0.219 to 0.055 (W = 0, p = 0.031), the anti-correlation of dissimilarly tuned pairs vanishing, and the residual contrast (though present in individual animals) not established across the group (W = 3, p = 0.156).

To further resolve the dissociation between SUB and VIS, we compared the full correlation matrices across conditions (Figure 5C-D). When comparing correlations between room or stripe conditions, the two regions were indistinguishable: similarity of correlation structure between the two room exposures, or between the two stripe placements, was equally high in SUB and VIS (pooled same-context similarity, tuned cells: 0.848 ± 0.026 vs 0.838 ± 0.054; C1 → C7: W = 10, p = 1.0). Across a context change, on the other hand, the regions diverged in every animal. From room to darkness, pairwise correlations in the SUB structure were largely preserved (0.774 ± 0.039) while correlations in the VIS structure collapsed (0.324 ± 0.138; SUB vs VIS: W = 0, n = 6, p = 0.031, r = 1.00), with the same divergence from room to stripe (0.849 ± 0.025 vs 0.578 ± 0.086; W = 0, p = 0.031). Same-context similarity exceeded cross-context similarity in the VIS in every animal (W = 0, p = 0.031 for tuned and all cells), whereas the corresponding difference in the SUB was small and, while consistent in direction in five of six animals, at the floor of what n = 6 can establish (tuned cells W = 0, p = 0.031, driven by one near-zero animal, and all cells W = 1, p = 0.063). The regional divergence is not a difference in measurement quality, with split-half reliability of the correlation matrices exceeding 0.75 in every group and condition, the regional difference surviving correction for attenuation (W = 0, p = 0.031), and the ordering holding at every bin width from 20 ms to 1 s (Supp. Fig. 5-2). The subicular network therefore sustains its internal coupling structure across sensory contexts, while visual cortical structure is bound to the context that created it.

### Manifold characterization

Having established that contextual identity is reflected in the similarity structure of population activity, we next asked how these relationships are organized at the level of the population manifold. We therefore examined the geometry of population activity across the two regions and conditions.

The overall pattern of population activity distinguished the two regions’ representations geometrically (Figure 6A-B; PCA projections are used for visualisation only and carry no decoding load). In the cue-rich room (C1), the two dimensional projection of subicular population activity formed a ring-shaped manifold, with position along the manifold corresponding to the mouse’s heading, consistent with an attractor-like representation (Figure 6A, left). Under the single stripe (C2; Figure 6A, middle), the ring became anisotropically distorted with non-uniform density along it. In darkness (C5; Figure 6A, right), the ordering of population states by head direction was largely lost and no ring-like organisation was demonstrable. Visual cortical activity showed a similar but weaker geometric progression (Figure 6B), forming a ring-like arrangement in the room with less uniform angular coverage than in SUB, and collapsing onto a cue-dominated axis oriented toward the stripe and its diametrically opposite “shadow” direction under the single cue. However, the directional information could still be extracted though it concentrated along a single stimulus-defined dimension.

### Comparative population decoding across environments

We next asked whether the differences in population geometry across environments were accompanied by differences in the information that could be extracted from population activity. To this end, we decoded head direction and angular velocity from population activity using a nonlinear decoder operating directly on the smoothed spike space (Figure 6C).

The decoder was able to reconstruct the mouse’s head direction with high accuracy in the cue-rich room condition (C1), with median circular absolute decoding errors of 18.2° ± 1.4° in the subicular region and 20.3° ± 1.1° in the visual region (Figure 6D). Even though the geometric organization of population activity was partially degraded in the stripe conditions (C2-4, C6), decoding performance remained good for both regions, with the visual region performing better (C2: VIS 40.7° ± 2.8°; SUB 55.1° ± 5.3°, paired p < 0.001). Including untuned neurons improved decoding slightly in all conditions, especially in the stripe environment. In the dark (C5), decoding errors were close to chance levels (78°-81°; Figure 6D) in both regions. Overall, these results show that both SUB and VIS populations represent head direction at the population level. They differ in their geometries and their performances, and in their dependency on the visual environment.

Comparison of decoding of the angular velocity (Figure 6E) revealed significantly better performance in VIS than in SUB, across all conditions. On average, angular velocity decoding errors were moderate in room conditions C1 and C7, close to chance in stripe conditions in SUB and slightly better in VIS. Similar values were obtained in darkness (Figure 6C). Including all cells systematically improved velocity decoding both in SUB and even more so in VIS, indicating that the cells selected for HD tuning alone do not optimally cover velocity information. When the decoding model included all recorded neurons, regardless of tuning, both SUB and VIS showed an improved decoding accuracy, mainly in the stripe conditions.

### Cue-anchored bias and geometric warping of the directional code

We next examined how decoding errors were distributed across head directions. To assess whether the stripe related biases reflected systematic geometric distortion, we analyzed signed decoding errors as a function of angular distance from the visual cue (Figure 7). During all stripe session trials, both SUB and VIS showed a wave-like relationship between signed error and stripe orientation. The size of this effect was greater in amplitude for VIS and smaller for SUB (stripe session 1: 50.0° ± 5.2°; session 3: 44.8° ± 3.1°; all p < 0.01 ; Figure 7). Errors were minimal when the mouse was facing the stripe (binocular field, 0° error), returning a wave-like warping pattern centered on the visual reference. Training decoders on room activity (C1) and testing them in stripe condition (C2) reproduced the same modulation (mean amplitude ≈ 40°-45° across sessions), showing that the linear mapping between neural activity and heading remains valid across contexts but becomes systematically distorted relative to the landmark.

These findings reveal that head-direction representations in both subicular and visual populations are geometrically warped by salient cues. The SUB tends to retain an internally referenced manifold that remains relatively stable across sensory changes, while VIS rests on a cue-anchored reference frame. This complementary warping provides a neural mechanism by which internally generated orientation signals align with external visual structure. Together, these results demonstrate that both regions sustain internally coherent population organization, yet differ in their reliance on external cues. The subicular network maintains ordered correlation structure across sensory transitions, consistent with an internally stable attractor framework, whereas the visual cortex exhibits cue-dependent coherence that weakens once landmark information is absent.

## Discussion

Spatial orientation emerges from the continual interaction of self-sustained dynamics with external landmark information. Here, we obtained the first simultaneous comparison of population activity in the subicular complex and the visual cortex during visual cue manipulation. We found that both regions carried directional signals and that both signals anchored to the visible landmark, as shifting the single cue shifted each region’s decodable directional frame by the corresponding angle. What distinguished the regions was the precision and stimulus-dependence of the readout, with the VIS decoding as accurate as the SUB decoding in the cue-rich room, but more accurate and sharper when exposed to a single high-contrast stripe, as can be expected of a retina derived signal. In darkness, the SUB pairwise coupling structure persisted while the VIS structure collapsed.

### Subicular dynamics and the persistence of internal coherence

We demonstrate that head-direction signals of single units in the subicular complex in passively rotated mice follow the classical definition of head-direction cell firing in the pre- and postsubiculum (Js Taube et al. 1990a, 1990b; Tukker et al. 2015). Tuning curves were stable over time, they covered the 360° of an environment uniformly, and preferred directions of firing followed the rotation of a visual cue, replicating canonical observations in freely moving animals (Figures 2-4; Goodridge et al. 1998; Knierim and Hamilton 2011). The defining characteristics of HD coding thus persist in the subicular network even in the absence of active locomotion, consistent with the preservation of HD firing under passive movement in the ADn (Shinder and Taube 2011). On the population level, we show that internal correlation coefficients amongst similarly tuned neurons remained high across different conditions and the pairwise correlation structure retained the ordering imposed by directional tuning (Figure 5; Supp. 5-1), the same population property that survives loss of anchoring in attractor models and thalamic HD ensembles across brain states and across changing reference frames (K. Zhang 1996; Peyrache et al. 2015; Chaudhuri et al. 2019; Park et al 2019; Yoganarasimha et al 2006). These observations verify that the subicular directional network retains its internal organization in darkness, in accordance with theoretical models of ring attractors (Skaggs et al. 1995; K. Zhang 1996; Peyrache et al. 2015; Simonnet et al. 2017; Laurens and Angelaki 2018), and with the months long preservation of the internal organization of the HD system (Skromne Carrasco et al. 2026).

The apparent loss of direction tuning in darkness (C5) is therefore not a loss of directional activity but an unanchoring of its internal representation. If the self-generated bump of population activity drifts, the average firing rate maps appear untuned, while the structure of the correlation and population embeddings still maintain their ordered topology. This interpretation fits with theoretical predictions and previous reports that the HD activity persists and drifts in cases of sensory deprivation (Js Taube 1995; Jeremy P. Goodridge and Taube 1995; Valerio and Taube 2012; Peyrache et al. 2015; Asumbisa et al. 2022). We also note that what survived cue removal at the single cell level was angular velocity and not direction, with the fraction of velocity-tuned cells rising in darkness even as directional tuning collapsed, consistent with an intact vestibular drive updating an unanchored integrator (Valerio and Taube 2012; Laurens and Angelaki 2018). The continual persistence of internal coherence under these conditions indicates that the subicular network acts as a self-sustaining substrate of orientation.

The restoration of visual landmarks reinstated the original directional maps (C1 vs C7), showing that the subicular ensembles can reset their internal dynamics to fit in with their structural associations with the environment when stable references are made available again (J. P. Goodridge et al. 1998; Zugaro et al. 2003; Yoder et al. 2015; Skromne Carrasco et al. 2026). The residual mis-alignment after the dark period (17 ± 6° against the reinstated stripe, versus 2 ± 1° in visual cortex; Figure 3B) measures the drift the internal estimate accumulated while unanchored. This pattern is consistent with landmark control and attractor dynamics in the HD system, where subicular circuits preserve not only the instantaneous heading but also inter-relational geometry between preferred directions, and thus allow the network to re-anchor its configuration when the sensory frame of reference returns (J. P. Goodridge et al. 1998; K. Zhang 1996; Skaggs et al. 1995; Peyrache et al. 2015; Chaudhuri et al. 2019). Together our work shows that the population activity in the subicular complex preserves the structural coherence of the head-direction attractor whilst displaying flexibility in its alignment to the external sensory frame of references.

### Visual cortical directionality: precision and dependence on sensory anchors

Directional coding in the visual cortex resembled head direction coding only at the first glance, and under our paradigm we were able to take apart this resemblance. With the platform rotating in the presence of a single fixed stripe, the stripe sweeps through the visual field, and the mouse’s binocular zone, straight ahead. A visual cortical neuron with an apparent PFD towards the stripe is in these conditions indistinguishable from a neuron responding as the stripe crosses its receptive field. We found that the data behaved accordingly, with the VIS tuning sharpened when the scene was reduced and its PFDs were pinned to the stripe, while its response at the stripe-free “shadow” direction was near zero and in the absence of visual input its coupling structure collapsed. In the room condition, with many landmarks tiling the panorama that drive responses across all angles, the ring-like organisation then is the multi-cue limit of the same mechanism. On this reading, the visual cortex does not itself compute head direction, but rather provides a precise landmark-bound directional reference.

That reference, however, is not merely reactive. The VIS populations carried more angular-velocity information than the SUB populations and, at matched cell counts in the room and in every condition including darkness, joining evidence that V1 receives head-motion signals of vestibular origins (Vélez-Fort et al. 2018; Guitchounts et al. 2020) and that cortical circuits merge visual and vestibular estimates of head velocity (Hennestad et al. 2021; Keshavarzi et al. 2022, 2023). Furthermore, directional and navigational variables are increasingly found in the visual cortex aligned with hippocampal signals (Saleem et al. 2013; Mertens et al. 2023; Siegenthaler et al. 2025). Here, our contribution is to separate, under parametric cue control, which parts of that signal are stimulus bound (directional preference, precision and coherence) and which are not (velocity signal and residual ensemble coupling). The visual system thus operates as an adaptive and calibrating structure for the HD system, dominating the anchoring of the compass when landmarks are informative and relinquishing control in their absence. Our finding of population level symmetry in the SUB with a single stripe should not be taken to mean an absence of cue aligned modulation at the cell level. Cells firing toward the stripe were modulated in SUB too (Figure 4F), and object based studies in freely moving mice have reported asymmetric facilitation/suppression (Siegenthaler et al. 2025), that is consistent with differences in cue type, behavioural state, and the effects of population averaging.

We find two complementary operating modes to exist within the distributed HD system. The subicular complex provides the coherent relational scaffold of the compass while the visual cortex, on the other hand, provides the precise cue-locked alignment of that scaffold to the external world. In our experimental conditions these two modes of operation become separable as the landmark can be removed or rotated. Landmark removal leaves the subicular scaffold intact while it abolishes the visual reference. Landmark rotations re-anchor both to the new placement. In this way, the neural compass combines internally generated stability with external correction, providing a continuous sense of heading across sensory environments.

### Integration and complementary coding across regions

Cross-region comparison of decoding performance highlights the complementarity of subicular and visual populations. Under cue rich room conditions (C1 and C7), the two regions were statistically indistinguishable (13.4 ± 1.8° vs. 12.2 ± 1.1°), while under a single stripe (C2-C4 and C6), the visual cortex was more accurate in every condition when the subicular error roughly tripled. Note that it is not evidence against a subicular compass. Within stripe blocks, part of the subicular error is not noise but a coherent and cue-locked deformation of the decoded frame, with 22.3 ± 4.8° peak-to-trough under the stripe against 7.1 ± 1.3° in the room, exceeding each animal’s own circular-shift null in six of six animals (Figure 7). The subicular representation re-anchors to any available landmark and inherits that landmark’s distortion, while the visual cortical representation is locked to the visual stimulus and is therefore most precise exactly when a single unambiguous landmark defines the scene. Furthermore, the visual cortex also carried more angular velocity information than SUB in every condition, including darkness (significantly at matched cell counts; Figure 6E). When all recorded neurons were included, the decoding improved in both regions and in every condition, indicating that directional and velocity information is distributed beyond the classically tuned subpopulations. Combining the regions, by contrast, added nothing once cell count was matched (not shown), the joint population never outperforming the single region, so the two directional readouts are largely redundant, as an inherited common anchoring signal would predict and as the shared deformation shows at the population level (Figure 7). The differences between SUB and VIS therefore reflect not the presence vs. absence of cue anchoring, but rather the relative weighting of an internally maintained and re-anchorable estimate against an externally defined reference (Angelaki and Laurens 2020; Keshavarzi et al. 2023).

### Population geometry and correlation structure

Low dimensional analysis showed that under stable visual conditions, population activity in both areas was organized in a continuous circular manifold, the topology expected of a ring attractor. In the subicular complex, this manifold spanned the whole range of orientations found in the room, and retained a partially circular geometry in the stripe condition (C2; Figure 6A). When all visual structure was removed (dark, C5), the SUB manifold lost demonstrable ring organisation and heading could no longer be decoded above the chance floor. What then persisted was not the geometry of the embedding, but the coupling structure that generates it (Figure 5). VIS population activity, on the other hand, did not express a full ring in the room condition (C1; Figure 6B). In the stripe condition (C2), the manifold became elongated and cue-biased, and in darkness (C5) the organization was lost. We suggest that in the visual cortex, the geometric arrangement is a reflection of the panorama and, accordingly, does not outlive the visual stimuli.

Analysis of pairwise correlations demonstrated the attractor organization of directional signals in the subicular complex. Aligned subicular pairs remained more correlated than non-aligned pairs in every condition including darkness (Δρ 0.11–0.20; W = 0, p = 0.031 at each condition), and the tuning-ordered correlation gradient retained 75 % of its span without visual input. Visual cortical coherence was equally structured under stable cues but collapsed across context changes (similarity 0.84 within context versus 0.32 from room to darkness), and its residual alignment contrast in darkness was weak and inconsistent across animals. This dissociation suggests that the SUB correlations reflect an internally stabilized attractor which maintains its functional topology, across sensory states, whereas VIS coherence depends largely on the external structure and decays rapidly when it disappears.

### A single anchoring signal read at two levels

In the final figure of this study (Figure 7), we show the deformation that a single landmark imposes on the decoded direction in our head-fixed paradigm. The deformations are cue-locked across SUB and VIS in every animal. We suggest that this similarity comes from inheritance of the anchoring signal. First, the observed deformation is caused by the cue in both regions, as it exceeded each animal’s own circular-shift null and collapsed in the multi-landmark room. Second, the subicular deformation is an attenuated copy half the amplitude of the visual cortical one in every animal, as expected from the combination of a biased visual anchoring input with an unbiased vestibular estimate of the platform’s motion. Third, we find that the same compromise appears independently at different timescale: after a 90° cue rotation the subicular map under-rotated to +65°, again landing between the visual frame and the vestibular one (Figure 3C). While the visual anchor moves with the stripe, the SUB first harmonic is too small in half the animals for its phase to be estimated, and the difference between the two anchors is unresolved. Taking account of these features, this can be explained by a visually derived anchoring signal reaching the subicular compass, a solution anatomically plausible via retrosplenial input to the presubiculum, where thalamic and retrosplenial signals converge on single L3 neurons (Yoder et al. 2011; Jacob et al. 2017; Richevaux et al. 2025).

Two further observations constrain how landmark information enters the directional circuit. First, the anchoring is both rigid and global as, after a cue rotation, decoders trained under the previous placement read out heading rotated by exactly the imposed angle, in both regions and every animal. The landmark therefore sets the phase of an otherwise preserved mapping rather than reshaping it. Second, the influence of the cue is not confined to the cue’s own direction, with SUB cells preferring the “shadow” direction retaining 39 % of the population’s cue-directed response against 16 % in VIS (Figure 4F). This mirror-axis structure may reflect the rotational symmetry of the ring-attractor connectivity, which propagates an anchoring signal applied at one direction around the entire ring (K. Zhang 1996; Knierim and Zhang 2012), and could be reinforced by bilateral visual pathways (Jacob et al. 2017; N. Zhang et al. 2022; Sit and Goard 2023). An alternative we cannot exclude is inference by elimination, since encoding that there is no stripe in view together with recent vestibular history can resolve directions away from the cue without symmetric connectivity. Still, the angular occupancy of cue and shadow directions was matched, excluding sampling artefacts, and the deformation collapsed in the room, placing the single cue as its cause. Landmark-based stabilisation thus extends beyond the cue itself, supporting the global continuity of the attractor while locally distorting its metric (Figure 7; Yoganarasimha et al. 2006).

Revealing that under sensory controlled conditions subicular and visual circuits carry a common directional signal read at two stages, our data allow the complementary roles of internal attractor dynamics and visual anchoring in orientation to be delineated. The subicular complex maintains the internal coupling structure of its directional network, which persists during sensory deprivation, even if the representation drifts, forming the basis for sustained orientation. The visual cortex injects context-specific precision and external congruence. Their interaction results in a compass having both stability and flexibility, continuity during sensory loss and recalibration as landmarks are reinstated. This interaction illustrates how distributed neural systems interface self-generated and sensory variables to maintain the stability of internal variables, a processing motif which may be relevant not only to navigation but to perception, memory, and decision making.

## Acknowledgements

This work was supported by the Centre National de la Recherche Scientifique and the Université Paris Cité. DF received funding from the Agence Nationale de la Recherche (ANR-DFG Program, ANR-18-CE92-0051 BURST), from the ERA-NET NEURON Program (ANR-20-NEUR-0005 VELOSO), and the FLAG-ERA HBP Program (ANR-21-HBPR-0002 VIPattract). This work has benefited from support by the BioMedTech Facilities at Université Paris Cité (Institut National de la Santé et de la Recherche Médicale Unité S36/Unité Mixte de Service 2009). We thank Fabrice Licata and Dr. Abdelali Jalil from the UPC Microscopy platform. We thank Mathieu Beraneck for discussions and for help with setting up experiments and advice on the visual projection.

## Author contributions

Experiments were designed by FA and DF. MG, IC and MW helped with setting up experiments. Electrophysiological recordings and histological analysis were carried out by FA. Data were preprocessed by FA, and analyzed by MD, including pipeline development, with the help of JL, MW and MG. MD, FA, MG and DF interpreted data. The manuscript was written by MD and DF with contributions from all authors.

